# Selective Androgen Receptor Modulator Microparticle Formulation Reverses Muscle Hyperalgesia in Mouse Model of Widespread Muscle Pain

**DOI:** 10.1101/2022.08.02.502353

**Authors:** Joseph B. Lesnak, David S. Nakhla, Ashley N. Plumb, Alexandra McMillan, Sanjib Saha, Nikesh Gupta, Yan Xu, Pornpoj Phruttiwanichakun, Lynn Rasmussen, David K. Meyerholz, Aliasger K. Salem, Kathleen A. Sluka

## Abstract

Currently, there is a need for the generation of non-opioid analgesics for treating chronic pain. Preclinical and clinical studies demonstrate the analgesic effects of testosterone. However, treatment with testosterone is not feasible due to adverse effects. Selective androgen receptor modulators (SARMs) were developed to overcome these limitations by minimizing activation of androgenic side effects. First, we demonstrate SARM administration alleviates widespread muscle pain in male and female mice. We then developed a SARM-loaded PLGA microparticle formulation that reverses widespread muscle pain in two injections. In vitro and in vivo release kinetics demonstrate the microparticle formulation had sustained SARM release for 4 weeks. Antagonism of androgen receptors blocked the analgesic effects of the SARM microparticles. SARM treatment had no effect on cardiac or liver enzymes, cardiac histology, and did not produce rewarding behavior. These studies demonstrate SARM microparticles as a potential therapeutic for chronic muscle pain.

**One Sentence Summary:** A selective androgen receptor modulator microparticle formulation alleviates widespread muscle pain in male and female mice while being non-toxic.

## INTRODUCTION

Musculoskeletal pain affects 13-47% of the population and costs the United States over $600 billion in health care costs and lost wages (*1*). Unfortunately, many current analgesics for chronic pain have negative side effects or are addictive; thus, there is an urgent need for the development of safer therapeutics for pain relief. While testosterone, which activates androgen receptors, is analgesic in both animal and human studies (*2-5*), there are significant unwanted effects. Medically, those taking testosterone require regular appointments with health care providers to monitor serum testosterone levels and adverse effects (*6*). Activation of androgenic pathways by testosterone results in negative side effects such as impaired fertility and virilization (*7*). Non-steroidal selective androgen receptor modulators (SARMs) have tissue specific anabolic effects, and limit unwanted androgenic effects (*8-10*). Preclinical and clinical work shows daily SARMs reduce bone loss, muscle atrophy, and fatigue associated with cancer treatment (*11-14*) with minimal side effects and are non-addictive (*8, 12, 15*). Thus, SARMs have potential to be therapeutic for individuals with chronic musculoskeletal pain.

Daily administration of drugs is associated with reduced adherence and consequent suboptimal dosing. Adherence to daily administration of prescribed medications is poor with roughly 50% of individuals with chronic diseases, including chronic pain, being non-adherent (*16-20*). It is estimated that medication noncompliance costs the United States $100 billion per year due to increased healthcare costs, increased morbidity and death (*21*). Prescription adherence decreases with daily dosing frequency of twice (Odds Ratio 1.2-4.8) and three daily administrations (Odds ratio 8.6) when compared with once daily dosing (*22-25*); while prescription adherence improves with once weekly administration (Odds ratio 1.6-1.9) when compared to daily dosing (*26, 27*). Further, repeated oral administration leads to fluctuations in drug plasma levels resulting in sub-therapeutic or toxic drug concentrations and consequent treatment failure or unwanted adverse effects (*17, 19, 20, 28*). One way to overcome poor adherence and fluctuating drug concentrations of orally delivered drugs is with long-acting, injectable microparticle formulations. These formulations allow for controlled drug delivery by slowly releasing drug over time to provide steady plasma levels of drug following a single administration (*29-31*). Poly (lactic acid-co-glycolic acid) (PLGA) is a synthetic, biocompatible and biodegradable polymer used to encapsulate drugs into microparticles to provide long-term drug release (*32*). Thus, PLGA microparticles increase adherence and ensure drug concentrations remain within a therapeutic window to improve treatment success. Therefore, we hypothesized that a SARM-loaded PLGA microparticle formulation would alleviate muscle hyperalgesia in a mouse model of widespread muscle pain.

## RESULTS

### Daily SARM Administration Alleviates Muscle Hyperalgesia

To determine if SARM administration alleviates muscle and paw hyperalgesia, we first tested if daily, systemic administration of SARMs reduced pain behaviors in mice. We utilized a widespread muscle pain model produced by two intramuscular acidic saline injections into the left gastrocnemius muscle which results in bilateral muscle and paw hyperalgesia (*33*). After induction of the pain model, daily administration of SARM (25 mg/kg, s.c.) reversed the decreased muscle withdrawal threshold (MWT) on both the ipsilateral (GroupXTime effect: F_5,65_= 3.21, p=0.01, Cohen’s *f*=0.50) and contralateral (GroupXTime effect: F_5,65_=4.12, p<0.01, Cohen’s *f*=0.56) side when compared to animals receiving vehicle injections (Fig. 1A+B). Analysis of area under the curve (AUC) showed that mice receiving SARM had a greater reduction in MWT when compared with animals receiving vehicle treatment (Ipsilateral p=0.01; Contralateral p=0.03) (Fig. 1C). However, SARM administration had no effect on mechanical sensitivity of the paw on either the ipsilateral (GroupXTime effect: F_5,65_=1.31, p=0.27, Cohen’s *f*=0.32) or contralateral (GroupXTime effect: F_5,65_=1.68, p=0.15, Cohen’s *f*=0.36) side when compared to animals receiving vehicle injections (Supplemental Fig. 1A+B). Thus, SARM alleviates muscle hyperalgesia in an animal model of widespread muscle pain.

**Figure 1.**
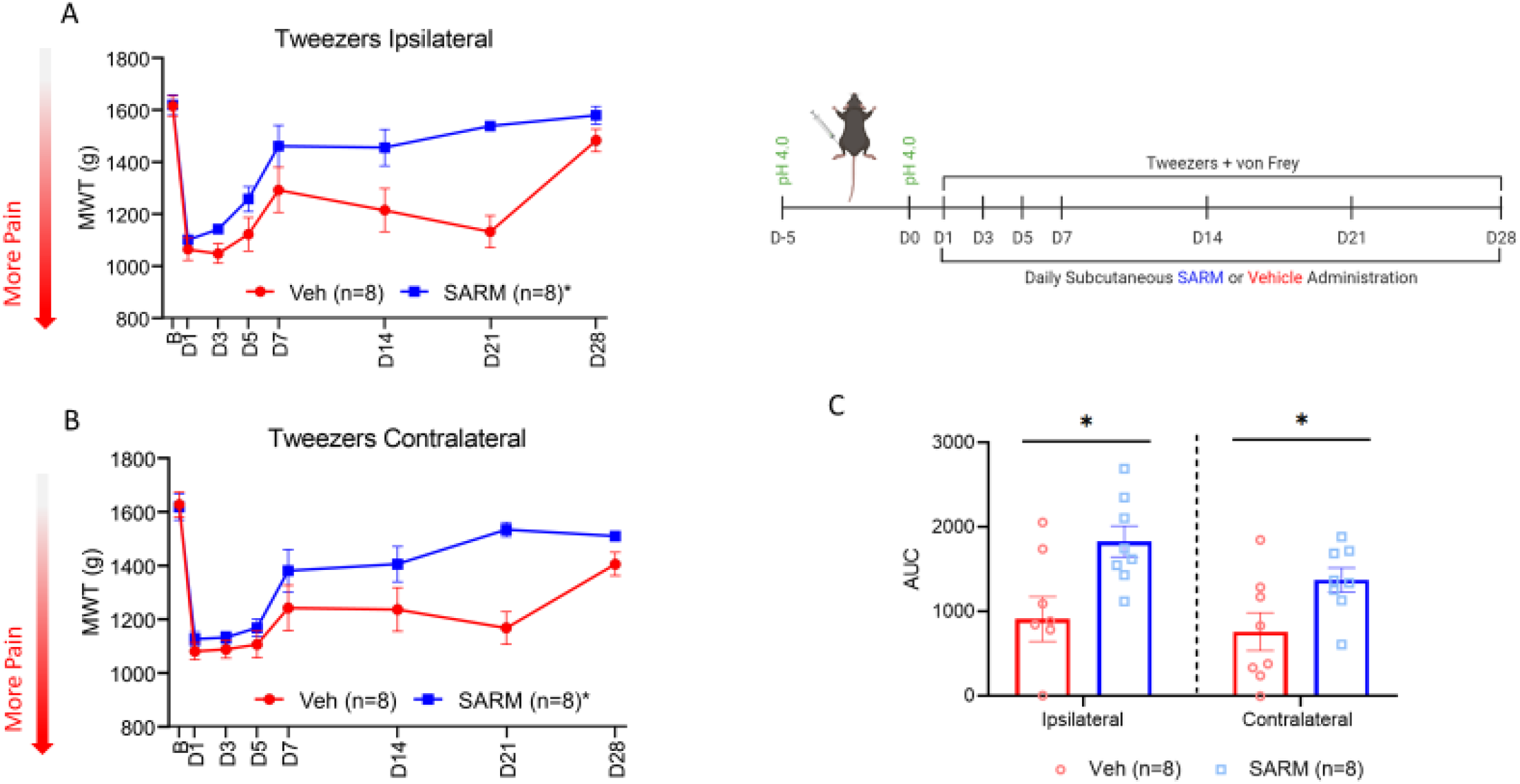
Daily SARM injection reverses muscle hyperalgesia. Following induction of the acidic saline muscle pain model, animals received daily subcutaneous injection of SARM (25mg/kg) or its vehicle for 4 weeks while pain measures were reassessed on days 1, 3, 5, 7, 14, 21, and 28. (A+B) Daily SARM administration reversed muscle hyperalgesia measured via muscle withdrawal threshold (MWT) on both the ipsilateral and contralateral limb. (C) Area under the curve analysis (AUC) demonstrates animals receiving SARM had a quicker recovery of MWT values than those receiving vehicle. *p<0.05 compared with vehicle; D=day, B=baseline, Veh=vehicle; Data are mean±SEM; Images made on Biorender.

### Development of PLGA SARM Microparticle Formulation

To overcome adherence issues with daily administered drugs reported in clinical studies (*16-20*), we developed a slow-release microparticle formulation for delivery of the SARM over a 4-week period. To determine the optimum formulation, we developed two different sized SARM-loaded PLGA microparticle formulations (Average diameter ± SD = 47.7 µm ± 3.4 and 3.6 µm ± 0.38 for F1 and F2, respectively) (Fig 2A). SARM-loaded PLGA microparticles (F1 and F2) had significantly different particle sizes (p<0.01, n=100) (Fig. 2B) and both formulations showed a homogenous, unimodal distribution with a narrow spread (Fig. 2C).

**Figure 2.**
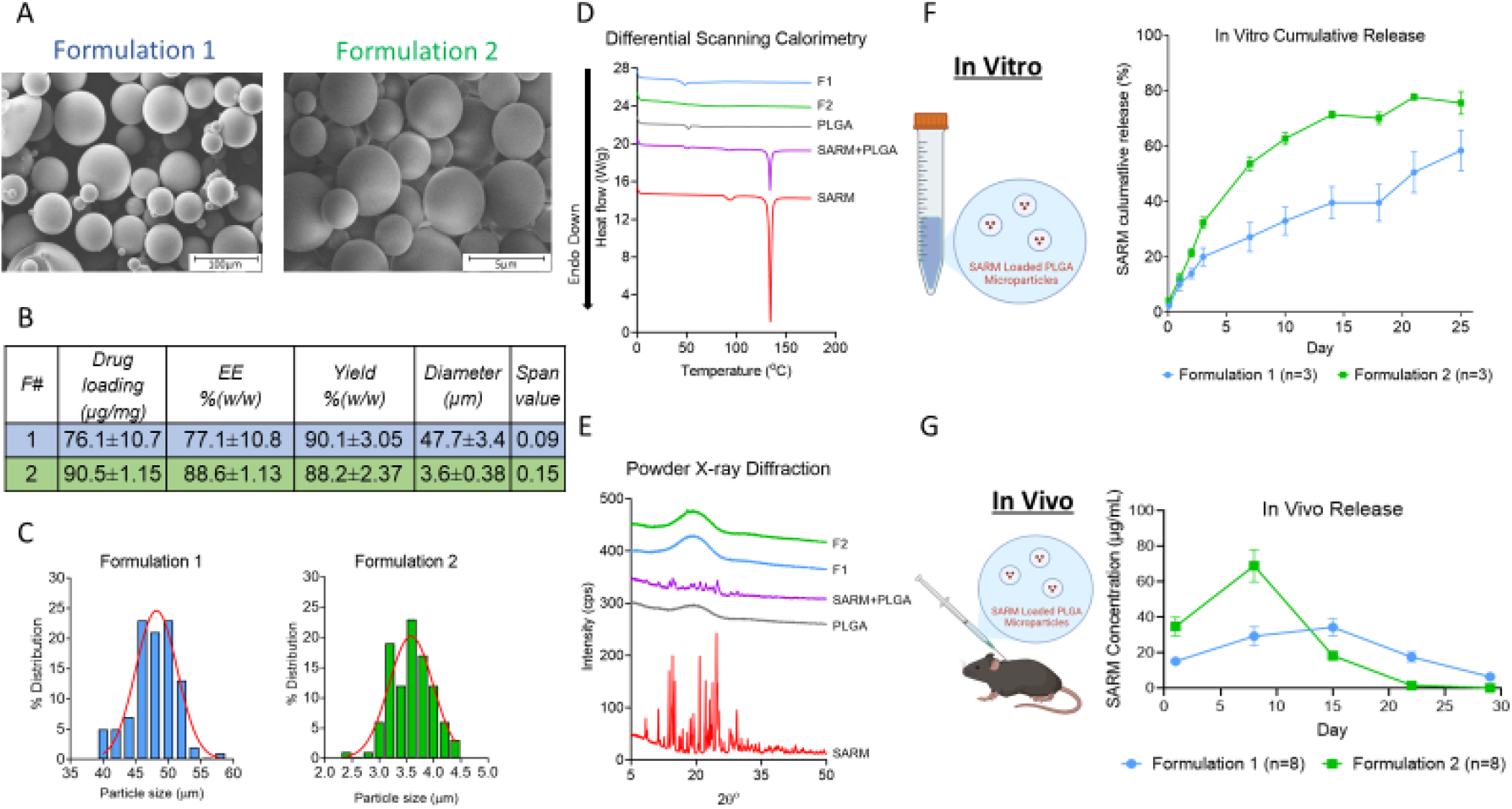
Development of SARM-loaded PLGA microparticle formulations. (A) Scanning electron microscope photomicrographs of SARM-loaded PLGA microparticles for formulation 1 and 2. The particles are spherical with a smooth, non-porous surface and no observable un-encapsulated drug crystals. (B) Table depicting drug loading, encapsulation efficiency, percent yield, and average diameter of the two microparticle formulations. (C) Particle size distribution histograms of the SARM-loaded PLGA microparticles formulations showing a unimodal, gaussian distribution indicating homogenous particle size distribution. (D) Differential scanning calorimetry thermograms of SARM microparticle formulation 2, SARM microparticle formulation 1, PLGA RG503H, mixture of SARM and PLGA RG503H, and pure SARM. (E) Powder X-ray diffraction diffractograms of SARM microparticle formulation 2, SARM microparticle formulation 1, mixture of SARM and PLGA RG503H, PLGA RG503H, and pure SARM. (F) In vitro cumulative release profiles of the SARM-loaded PLGA microparticles for formulations 1 and 2. (G) Mean plasma concentration-time profile of SARM following subcutaneous administration of SARM-loaded PLGA microparticles formulations at day 0 and day 7 at a dose of 60mg. F=formulation MP=Blank Microparticles; Data is presented as mean±SD for B and mean±SEM in F and G; Images made on Biorender.

To ensure the drug was encapsulated within the microparticle formulation, we used differential scanning calorimetry (DSC) and powder x-ray diffraction (PXRD). DSC tests if there is an interaction between the polymer and the encapsulated drug by evaluating for a change in the glass transition temperature (*Tg*) of the polymer or the melting point of the encapsulated drug (*34, 35*). The SARM-loaded PLGA microparticles do not show the endothermic peak observed with SARM alone (red) or with a mixture of PLGA and SARM (purple)(Melting point, 134.18^°^C; range 127.74 – 138.61^°^C) suggesting the SARM is encapsulated within the microparticle (Fig. 2D). Powder X-ray diffraction is used to determine whether a compound is in a crystalline or an amorphous state. When a crystalline drug is encapsulated in a microparticle formulation it loses its crystalline properties and exists in a molecularly dispersed (amorphous) form (*36*). The diffraction patterns of the SARM-loaded PLGA microparticles (F1 and F2) (Fig. 2E) show complete disappearance of the sharp peaks that are characteristic of the SARM crystalline state (red) indicating that SARM was encapsulated into the PLGA microparticles.

### SARM-loaded PLGA Microparticle Release Kinetics

To determine release kinetics of the two SARM-loaded PLGA microparticle formulations, we measured the cumulative release of encapsulated SARM over 25 days in an in vitro preparation. SARM-loaded microparticle formulations showed a typical bi-phasic release pattern characteristic of PLGA formulations (Fig. 2F). The cumulative release of SARM from F2 was faster than that of F1. After 3 days, 32.3% (±3.5) of SARM was released from F2 compared with 19.9% (±5.6) from F1. This was followed by a slow-release phase between days 3 and 18 and a faster release phase between days 18 and 25 for both F1 and F2. By day 25, F2 had released 75.5% (±7.05) of the encapsulated SARM while 58% (±12.6) had been released from F1.

Next, we analyzed SARM-loaded PLGA microparticle release kinetics in mice by measuring SARM’s plasma concentrations over time following injection of F1 and F2 (Fig. 2G). F1 showed a slower, more sustained release of SARM from microparticles with plasma levels detectable through Day 29 when compared to F2. Plasma levels for F1 ranged between 2.4 – 55.7 µg/mL which is within a therapeutically relevant concentration when compared to SARM plasma levels observed after IV or PO administration (*37*). In contrast, plasma levels of F2 SARM-loaded microparticles peaked at Day 8 and were not detected after Day 22.

### SARM-loaded Microparticles Alleviate Muscle Hyperalgesia

Based on the longer sustained release profile of F1, we tested if this formulation of SARM-loaded PLGA microparticles could alleviate muscle hyperalgesia. Treatment with F1 SARM-loaded PLGA microparticles significantly reduced muscle hyperalgesia bilaterally after induction of the muscle pain model when compared to vehicle treated animals (GroupXTime effect: Ipsilateral, F_15,100_=3.88, p<0.01, Cohen’s *f*=0.76; Contralateral, F_15,100_=5.12, p<0.01, Cohen’s *f*=0.88) (Fig. 3A). Mice that received two injections of SARM-loaded PLGA microparticles (days 1 and 7) showed a significant reduction in MWT when compared to vehicle treatment (Ipsilateral, p<0.01; Contralateral, p<0.01). No differences in MWT were found, compared to vehicle or two injections of SARM, for those that received a single injection of SARM-loaded PLGA microparticles (day 1 or day 7) (p=0.07-0.94). Similarly, analysis of AUC confirmed that mice receiving two injections of SARM-loaded PLGA microparticles had a greater reduction in MWT when compared with mice receiving vehicle (Group effect: Ipsilateral F_3,23_=4.98, p=0.01, Tukey post hoc p<0.01 vs vehicle; Contralateral F_3,23_=5.61, p<0.01, Tukey post hoc p=0.01 vs vehicle) (Fig. 3B). Similar to daily SARM administration, SARM-loaded PLGA microparticles had no effect on mechanical sensitivity of the paw (GroupXTime effect: Ipsilateral F_15,100_=1.47, p=0.13, Cohen’s *f*=0.47; Contralateral F_15,100_=1.28, p=0.23, Cohen’s *f*=0.43) (Supplemental Fig. 1C+D). These data demonstrate that SARM-loaded PLGA microparticles reverse muscle hyperalgesia following two injections.

**Figure 3.**
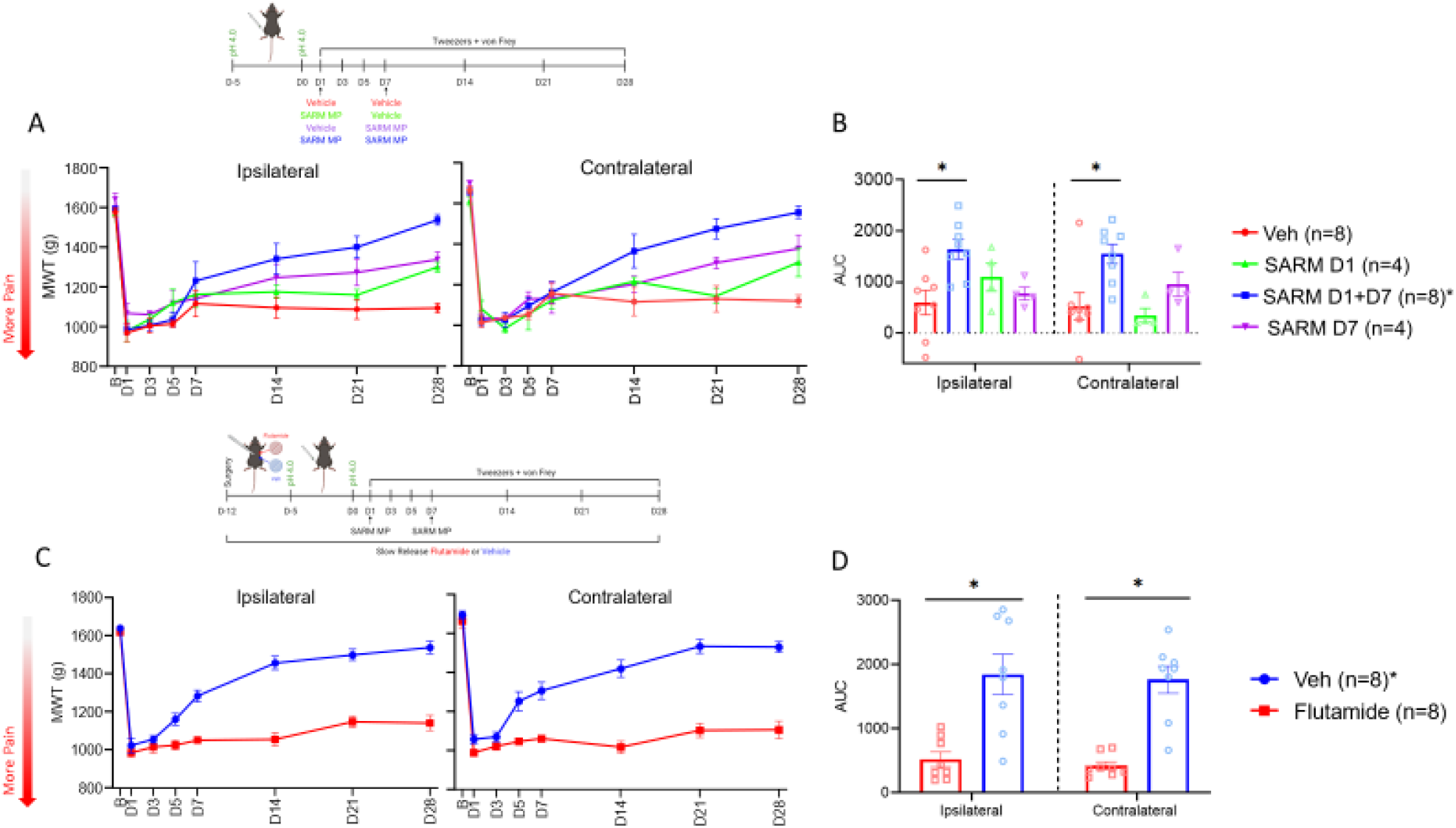
SARM-loaded PLGA microparticles alleviates muscle hyperalgesia and works through activation of androgen receptors. Following induction of the acidic saline pain model, animals received subcutaneous injection of SARM-loaded PLGA microparticles (60mg) on day 1, day 7, or on days 1 and 7 or received vehicle injections. Pain measures were reassessed on days 1, 3, 5, 7, 14, 21, and 28. (A) Animals who received SARM-loaded microparticles (Formulation 1) on days 1 and 7 saw a reversal of muscle hypersensitivity measured via muscle withdrawal threshold (MWT) on both the ipsilateral and contralateral limb when compared to animals receiving vehicle injections. There was no effect on MWT seen for animals who received SARM-loaded PLGA microparticles solely on days 1 or 7. (B) Area under the curve analysis (AUC) demonstrated animals receiving two injections of SARM microparticles had a quicker recovery of MWT compared with those receiving vehicle injections. In a separate experiment animals received implantation of a slow release flutamide (200mg) or vehicle pellets 7 days prior to induction of the acidic saline muscle pain model. Following induction of the pain model, animals received injections of SARM-loaded PLGA microparticles (60mg) on days 1 and 7 and pain measures were reassessed on days 1, 3, 5, 7, 14, 21, and 28. (C) Flutamide administration prevented the analgesic effects of the SARM-loaded PLGA microparticles on both the ipsilateral and contralateral limb measured via reversal of MWT decrease as compared with animals who received vehicle pellets. (D) AUC analysis revealed SARM’s ability to recovery MWT values was blocked by the implantation of flutamide pellets. *p<0.05 compared with vehicle; D=day, B=baseline, Veh=vehicle, MP=microparticles; Data are mean±SEM. Images made on Biorender.

### SARM-loaded Microparticles Alleviate Muscle Hyperalgesia Through Activation of Androgen Receptors

Since SARMs were developed to serve as androgen receptor agonists (*10*), we tested if SARM-loaded PLGA microparticles produce analgesia through activation of androgen receptors. To block androgen receptors during SARM treatment, we utilized slow-release pellets of the androgen receptor antagonist, flutamide. In the presence of flutamide, SARM loaded-PLGA microparticles did not reverse the muscle hyperalgesia when compared with animals implanted with vehicle pellets (GroupXTime effect: Ipsilateral F_5,65_=18.18, p<0.01, Cohen’s *f*=1.18; Contralateral F_5,65_=8.86, p<0.01, Cohen’s *f*=0.83) (Fig 3D). Similarly, analysis of AUC showed that mice receiving flutamide pellets with the SARM loaded-PLGA microparticles showed significantly less change in MWT when compared with animals who received vehicle pellets (Ipsilateral p<0.01; Contralateral p<0.01).

### SARM Toxicity and Rewarding Behavior

Due to clinically documented SARM induced alterations in liver enzymes (*11, 38-40*), we tested SARM-loaded PLGA microparticles for liver and heart toxicity. Liver and cardiac toxicity panels were run in mice 4 weeks after the first of two injections of SARM-loaded PLGA microparticles or its vehicle. There were no significant differences in levels of alkaline phosphatase, aspartate transaminase, albumin, total bilirubin, alanine aminotransferase, total protein, and creatine kinase between SARM microparticle-treated and vehicle-treated groups (p=0.02-0.82) (Fig. 4A). Microscopic examination of the heart tissue 4 weeks after SARM-loaded PLGA microparticle or vehicle treatment showed no differences between groups (Fig 4B). There were also no significant differences in body weights between animals receiving two injections of the SARM-loaded microparticles when compared with those receiving two vehicle injections (F_1,14_=1.19, p=0.29) (Fig. 4C). Since SARMs are also being used as a performance enhancing drug by athletes (*41-43*), we decided to test if regular SARM administration could be addictive. To do so, we tested if SARM produced rewarding-like behaviors using conditioned place preference testing. Following 5 days of pairing, animals did not show an increased preference for the SARM paired chamber between baseline (380.3±51.0sec) and post drug pairing testing (385.7±37.1sec) (t_7_ = 0.15, p=0.88) (Fig. 4D). Together, this data indicates that the SARM-loaded PLGA microparticles did not induce any cardiac or liver toxicity and that soluble SARM did not produce reward like behavior in mice.

**Figure 4.**
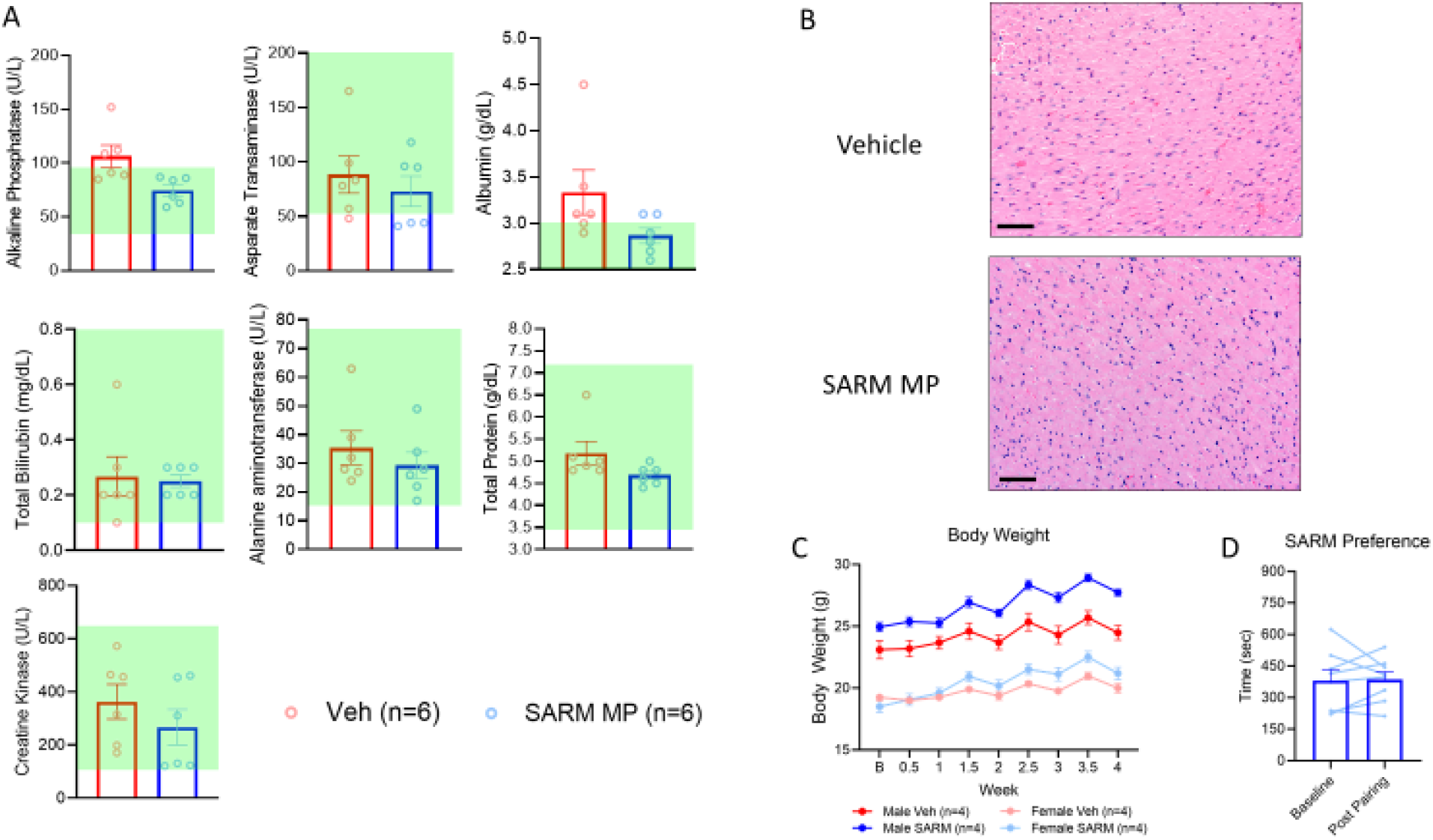
Safety and toxicology screening for animals who received 2 injections of SARM loaded PLGA microparticles. Serum and heart tissue was collected on day 28 from animals who received SARM loaded PLGA microparticles or vehicle injections both on days 1 and 7. (A) Serum levels of biomarkers for liver and cardiac toxicity showed no differences between animals receiving SARM loaded PLGA microparticles or its vehicle. Average serum levels also fell within reported normal ranges for each biomarker in animals receiving SARM-loaded microparticles as represented by green shaded boxes. (B) Histological (H&E staining) examination of the heart tissue following four weeks of treatment did not show differences between animals who received vehicle or SARM loaded PLGA microparticles. (C) Body weights measured throughout the 28 days of treatment revealed no differences between animals receiving SARM-loaded PLGA microparticles and vehicle injections. (D) Condition place preference testing revealed 5 days of SARM pairing (25mg/kg) produced no preference for the SARM paired chamber. Veh=vehicle, MP=microparticle; Black bar represents 65 μm; Data are mean±SEM

## DISCUSSION

The current study is the first to show that systemic administration of a SARM can be used to alleviate widespread muscle hyperalgesia using an animal model. We developed a SARM-loaded PLGA microparticle formulation that produced long-term release, *in vitro* and *in vivo*, and long-term reduction in hyperalgesia. The alleviation of hyperalgesia by the SARM microparticle formulation was prevented by flutamide, suggesting that the analgesic effect of SARMs works through activation of androgen receptors. There were no adverse changes in body weight, cardiac and liver toxicity enzymes, and heart histology in mice treated with SARM-microparticles suggesting that the formulation was safe. Thus, we developed a SARM-loaded slow release microparticle formulation that reduces hyperalgesia and is safe.

While we are the first to use SARMs to target androgen receptors for relief of muscle pain, this work is in agreement with preclinical and clinical studies showing analgesic effects of testosterone. In animals, testosterone reduces hyperalgesia in models of muscle, temporomandibular, formalin-induced, and stress-induced pain (*2-4, 44*). Clinically, women with fibromyalgia (n=12) treated with testosterone for 4 weeks showed a reduction in muscle pain, stiffness, and fatigue when compared to pre-treatment ratings (*5*). There are mixed results in hypogonadal or androgen-deficient men with some showing testosterone treatment reduces pain (*45, 46*), while others show no effect (*47*). While we demonstrate SARM administration reverses muscle hyperalgesia, it was unable to alleviate paw hyperalgesia. As the SARM used in the current study targets muscle and bone androgen receptors (*8, 48, 49*), it is possible that it is more effective for muscle pain than cutaneous pain. Alternatively, the SARM could target the site of insult over secondary sites; however, we see a reduction in the contralateral, uninjured muscle, suggesting the SARMs work to reduce hyperalgesia outside the site of insult.

We confirmed that SARM produced analgesia through activation of androgen receptors. Androgen receptors are located throughout the body, including both peripheral and central nociceptive sites, and on immune cells that modulate nociception (*50*). Since we administered the SARM systemically, the site of action of the SARM remains unclear. Activation of androgen receptors transcriptionally increases expression of mu-opioid and cannabinoid type-1 receptors on nociceptors, both of which have endogenous anti-nociceptive effects (*51-54*). Activation of androgen receptors also decreases pro-inflammatory cytokines (*55-58*) and increases anti-inflammatory cytokines (*56-60*) from a variety of immune cells, and can polarize resident macrophages to an M2 anti-inflammatory phenotype (*61, 62*). Prior studies show anti-inflammatory cytokines such as IL-10 and IL-4, are analgesic (*63, 64*). Lastly, lower amounts of circulating testosterone are associated with lower activation of the rostral ventromedial medulla, a brain region responsible for descending pain inhibition (*65*). Since, androgen receptors are located in a diverse range of tissues, it likely the analgesic effects of SARM are multifactorial.

SARMs have a half-life between 16 – 24 hours which suggests that daily administration is necessary to maintain therapeutic plasma concentrations that will reverse muscle pain (*66*). Daily administration results in lower adherence when compared to weekly or monthly administration (*16-18*) and could result in failure of SARMs to produce analgesic effects. For those with chronic diseases, long-term release of pharmaceutical agents can improve adherence, symptoms, and disease management (*67*). Several studies report non-adherence to prescribed medication in individuals with chronic pain and have indicated various reasons for non-adherence, including side effects, dosing frequency, poly-medication, and both high and low pain intensity (*17, 19, 20*). Injectable polymeric long-acting formulations are clinically available, safe, and have the potential to overcome adherence issues that result from repeated administration of medications (*29, 31, 67-69*). Thus, we developed a novel injectable, long-acting SARM-loaded PLGA microparticle formulation that can be injected subcutaneously to provide sustained release and maintain effective plasma levels of SARM over a long period of time, thus eliminating the need for daily administrations.

The current study demonstrates that a larger microparticle (Avg diameter = 47.7 µm) produces a more sustained release of SARM *in vitro* and *in vivo*. This particle size is similar to currently available compounds that are delivered subcutaneously through 20–21-gauge needles and include triptorelin pamoate, risperidone, and naloxone (*31*). The long-term release of our SARM-loaded PLGA microparticle formulation, at least 29 days, with accompanying analgesia suggests adequate plasma levels can be achieved with a single or twice per month administration. Thus, the microparticle formulation has the potential to produce long-term reduction in pain.

The lack of toxicity in the current study for SARMs agrees with prior work in both preclinical and clinical studies. The weights of the prostate and seminal vesicle are unchanged after the administration of SARMs to orchiectomized rats when compared to gonadally intact male rats (*13*). Similarly, in humans there are no adverse androgenic events such as virilization, hirsutism, or negative effects on the prostate as reported in randomized double-blind placebo controlled clinical trials in both males and females (*11*). Numerous clinical studies utilizing SARMs examined safety and found similar number of adverse events between treatment and placebo groups (*11, 38-40*). No serious adverse events were reported due to drug treatment (*11, 38-40*). Other adverse events reported include constipation, dyspepsia, nausea, muscle soreness, headaches, fatigue, increased hemoglobin and creatine kinase, and decreased high density lipoproteins (HDL) (*11, 38-40*). The most consistently cited adverse event with SARM treatment is an increase in the liver enzyme alanine-amino-transferase (ALT) (*11, 38-40*). However, we found no alterations in ALT in the current study in response to SARM-loaded microparticle treatment. Differences could be related to the microparticle formulation’s ability to maintain stable levels, a shorter duration of treatment, or species differences between the mice and humans. There are also published case reports of liver injury due to SARM misuse in young, healthy individuals (*41-43*). Since SARMs have anabolic effects that increase muscle growth and enhance athletic performance, they have become a drug of interest for athletes and weightlifters. In these case reports, the dose of SARM used was either unreported or 10-100 times higher than that used in clinical trials (*41-43*). Thus, SARMs should still be viewed as safe when taken at appropriate doses.

The current study suggests activation of androgen receptors through SARM as a new potential treatment of chronic muscle pain. Activation of androgen receptors by testosterone are analgesic in both pre-clinical and clinical research (*2, 3, 5*). However, SARMs appear to be better tolerated with fewer adverse side effects due to minimal activation of androgenic signaling (*11, 13*). We also demonstrated that SARM-loaded PLGA microparticles produce long-lasting release for up to 4 weeks. Microparticle-based drug delivery could be ideal for individuals with chronic pain and could improve patient convenience, increase adherence, and optimize maintenance of therapeutic concentrations for a variety of pharmaceuticals. Future work should develop SARM-loaded microparticle formulations for testing in human subjects with chronic pain.

## Limitations

There are several limitations to this study. First, we only tested the analgesic effects of one class of SARMs. The SARM we utilized was designed to target bone and skeletal muscle tissue (*8, 48, 49*). Several SARMs have been developed that serve as site specific androgen receptor agonists at other locations such as the nervous system (*70*). It is possible that other SARM formulations could also alleviate muscle pain, however our results cannot be generalized to these other compounds. Next, we only tested the analgesic effects of SARMs in a mouse model of widespread muscle pain. It is unknown if SARMs would be analgesic in other pain models such as neuropathic or inflammatory pain, and therefore future studies should test the analgesic effects of SARMs in other pain models. Lastly, we only developed and tested one SARM microparticle formulation for its analgesic effects. This formulation was effective, and we were able to achieve full reversal of hyperalgesia with two doses. However, it is possible that a different particle size or dose could produce a greater effect with a single injection.

## MATERIALS AND METHODS

### Study Design

This study was designed as a randomized, blinded, controlled laboratory experiment using an equal number of male and female C57BL/6J mice for each experiment. All behavioral experiments were done in the morning, the investigator was blinded to group and all animals were randomly assigned to groups using a random number generator software program in blocks of 4, stratified by sex. For each experiment, male and female mice were evenly distributed, and multiple replicates were utilized. For each behavioral experiment, preliminary data was utilized to calculate samples sizes with power set at 0.80 and a significance set at 0.05. There were no data that needed to be excluded and no outliers were found. The main goal of this study was to develop a SARM-loaded PLGA microparticle formulation that could alleviate muscle pain.

*Experiment 1* tested if daily SARM administration alleviated muscle and paw hyperalgesia. Mice received the acidic saline muscle pain model and pain behaviors were assessed to confirm presence of muscle and paw hyperalgesia 24 hours after induction of the model. Mice received daily administration of SARM (25mg/kg, s.c.; n=8) or its vehicle (n=8) for the next 28 days. Pain-behaviors were reassessed on days 3, 5, 7, 14, 21, and 28 after induction of the pain model. The primary endpoint for this experiment was reversal of muscle hyperalgesia. Secondary endpoint was reversal of paw hypersensitivity.

*Experiment 2* developed 2 formulations of SARM-loaded PLGA microparticles that differed based on size of microparticle. Altering the size of the microparticles allows for control of SARM release rate where smaller microparticles release drug quicker and larger particles release drug slower. In this experiment we characterized diameter, drug loading, encapsulation efficiency, differential scanning calorimetry, and powder X-ray diffraction of each formulation.

*Experiment 3* tested release kinetics of the SARM-loaded PLGA microparticles in both *in vitro* and *in vivo* settings. For the in vitro experiment, each formulation (n=3) was placed in a release medium and SARM concentration was measured by high performance liquid chromatography (HPLC) at the following time points: 2 hour and 1, 2, 3, 7, 10, 14, 18, 21, and 25 days. For the in vivo experiment, animals were allocated to receive two injections (Day 0 and Day 7) of 60 mg of SARM loaded PLGA microparticle formulation 1 (n=8) or formulation 2 (n=8). Blood was collected on days 1, 8, 15, 22, and 29 after the first SARM microparticle injection and SARM concentration in plasma was measured by HPLC with ultra-violet spectroscopy (HPLC-UV). The primary endpoints for these experiments were cumulative SARM release *in vitro* and mice plasma SARM concentration over time, *in vivo*.

*Experiment 4* tested the analgesic effects of formulation 1 of the SARM-loaded PLGA microparticles. Mice received the acidic saline muscle pain model and pain behaviors were assessed to confirm presence of muscle and paw hyperalgesia 24 hours after induction of the model. Mice received 60 mg of SARM microparticles (n=16) or its vehicle subcutaneously (n=8). SARM microparticles were delivered at 24 hours (n=4), 1 week (n=4), or at both 24 hours and 1 week (n=8) following induction of the pain model. Pain-behaviors were reassessed on days 3, 5, 7, 14, 21, and 28 after induction of the pain model. The primary endpoint for this experiment was reversal of muscle hyperalgesia measured. Secondary endpoint was reversal of paw hypersensitivity.

*Experiment 5* tested if androgen receptor mediated the analgesic effects of SARM-loaded microparticles. Mice were implanted with a slow-release pellet of the androgen receptor antagonist flutamide (200mg; n=8) or a vehicle pellet (n=8) 7 days prior to induction of the pain model. Mice received the acidic saline muscle pain model and muscle hyperalgesia was confirmed 24 hours after induction. Mice then received subcutaneous injection of 60mg of SARM-loaded PLGA microparticles at 24 hours and 1 week after induction of the pain model. Muscle hyperalgesia was reassessed on days 3, 5, 7, 14, 21, and 28 after induction of the pain model. The primary endpoint for this experiment was reversal of muscle hyperalgesia.

*Experiment 6* tested the safety and toxicity of SARMs. Serum was collected from mice in Experiment 4 at the end of behavior testing on day 28 from animals who received two injections of SARM-loaded microparticles (n=6) or its vehicle (n=6). The serum was analyzed using liver and cardiac toxicity panels. Whole hearts were also collected from animals who received SARM-loaded microparticles (n=2) or its vehicle (n=2) and were analyzed for signs of toxicity via hematoxylin and eosin (H&E) staining. Lastly, we tested if repeated administration of soluble SARMs produces rewarding behaviors in mice using a conditioned place preference paradigm. Mice (n=8) received 5 days of pairing with subcutaneous SARM injections (25mg/kg). To determine if rewarding behavior was present, the amount of time spent in SARM paired chambers was compared between baseline and post-drug pairing.

### Mice

All experiments were approved by the University of Iowa Animal Care and Use Committee and were performed in accordance with the National Institute of Health guidelines. A total of 80 C57BL/6J mice (40 male, 40 female) (20-30 g) (8 weeks of age) (Jackson Laboratories, Bar Harbor, ME, USA) were used in the described studies. All mice were housed 4 per cage on a 12-hour light-dark cycle with access to food and water *ad libitum* unless noted otherwise.

### Acidic Saline Muscle Pain

Widespread muscle pain was produced through two intramuscular (i.m.) injections of acidic saline 5 days apart as previously developed by our laboratory (*33*). On days 0 and 5, mice were anesthetized with 2-4% isoflurane and injected with 20 μl of pH 4.0±0.05 saline into the left gastrocnemius muscle. This model produces bilateral paw and muscle hyperalgesia which lasts for up to 4 weeks (*33, 71*).

### Behavioral Assessments

Muscle hyperalgesia was assessed as muscle withdrawal thresholds (MWT) by applying force sensitive tweezers to the left and right gastrocnemius muscle as previously described (*72, 73*). Mice were placed into a gardener’s glove and the gastrocnemius muscle was squeezed with a pair of custom-built force sensitive tweezers until the animal withdrew its limb or vocalized. Both the left and right gastrocnemius muscle were tested and an average of 3 trials, 5 minutes apart, was used to determine MWT for each limb. A decrease in MWT was interpreted as muscle hyperalgesia.

Paw hyperalgesia was measured by examining the response frequency to repeated applications of a 0.04g von Frey monofilament. Mice were placed inside individual small cages on top of a wire mesh and allowed to acclimate for 60 minutes prior to testing. The von Frey filament was applied to the left and right hind paw 5 times over 10 rounds, 5 minutes apart. The number of withdrawals per round was then averaged for each paw. An increase in the number of responses was interpreted as paw hyperalgesia.

Conditioned place preference (CPP) testing examined for rewarding behavior. CPP boxes used for testing were 27.5” by 8.25” divided into two 10.75” by 8.25” pairing chambers and a 4.75” by 8.25” middle chamber. The pairing chambers had different colored walls (solid black vs black and white stripe) and floor textures (smooth vs ridged). The middle chamber had solid black walls with smooth floors and was illuminated. Acclimation and baseline testing occurred over the first 3 days. Mice were placed in boxes with access to all 3 chambers for 15 minutes. On the third day, time spent in each box was recorded. Drug pairing then occurred for 5 days on days 4 through 8. Each day, mice were injected subcutaneously with SARM vehicle (80:20; PEG300:DMSO) and placed in a pairing chamber for 60 minutes. Mice were then returned to their home cages. Four hours later, mice were subcutaneously injected with SARM (25mg/kg) and were confined to the opposite pairing chamber for 60 minutes. Chamber pairings were counterbalanced amongst mice. Following pairing, on day 9, mice were placed in boxes with access to all 3 chambers for 15 minutes with time spent in each box recorded.

### Drug Administration

The SARM, (s)-3-(4-cyanophenoxy)-n-(4-cyano-3-(trifluoromethyl)phenyl)-2-hydroxy-2-methylpropanamide (Achemblock, San Francisco, CA, USA), was used in the following experiments. For subcutaneous SARM administration, the drug was dissolved in dimethyl sulfoxide (DMSO; Fisher Scientific, Waltham, MA, USA) at a concentration of 50mg/mL and then diluted in polyethylene glycol 300 (PEG300; Med Lab Supply, Pompano Beach, FL, USA) until a final solution of 10mg/mL in 80:20 PEG300:DMSO was achieved. Vehicle control for subcutaneous SARM was 80:20 PEG300:DMSO. For administration, mice were anesthetized with 2-4% isoflurane and SARM (25 mg/kg) or vehicle were delivered subcutaneously. This dose was chosen based on prior literature which found greatest effect at this dose (*12*). SARM solutions were prepared fresh at the beginning of each week.

For SARM-loaded PLGA microparticles, a 60mg dose was utilized which is equivalent to a soluble SARM dose of 200 mg/kg based on a mouse weighing 25 grams. For administration, mice were anesthetized with 2-4% isoflurane and mice were subcutaneously injected with 60 mg of SARM-loaded PLGA microparticles suspended in 400μl of Dulbecco’s phosphate-buffered saline (DPBS; Gibco, Waltham, MA, USA).

Slow release flutamide (200 mg/pellet, 60-day release; Innovative Research of America, Sarasota, FL, USA) or control pellets were implanted subcutaneously. For implantation, animals were deeply anesthetized with 2-4% isoflurane. The pellet was implanted through a small incision at the nape of the neck. The incision was stitched closed with synthetic non-absorbable monofilament silk sutures. To protect the surgical site, animals were placed in single housed cages for the remainder of the experiment. Animals were allowed 7 days to recover from surgery prior to induction of the pain model. This dose was chosen based on prior literature showing that it blocked the analgesic effects of activating androgen receptors (*74*).

### Preparation of SARM-loaded PLGA Microparticles

Due to the hydrophobic nature of SARM, a single (oil-in-water) emulsion solvent evaporation technique was used to prepare the SARM-loaded PLGA microparticles (Supplemental Fig. 2). SARM and PLGA (Resomer RG 503 H) were dissolved in dichloromethane (DCM) at a concentration of 10 mg/mL and 100 mg/mL, respectively, and 1 mL of each solution was mixed (10 mg SARM, 100 mg PLGA; 1:10, drug:polymer ratio). The mixed SARM/PLGA solution was added to an aqueous phase (30 mL; 1% w/v polyvinyl alcohol in Nanopure water; Barnstead Thermolyne Nanopure water purification system, Thermo Fisher,Waltham, MA) and the mixture was immediately emulsified using either a Talboys Model 101 overhead mixer at speed 2833 rpm for 4 minutes (formulation 1, F1) or an overhead homogenizer (Ultra-turrax T25 basic, Ika Works, Inc., Wilmington, NC) at speed 9500 rpm for 1 minute (formulation 2, F2). The emulsion was transferred to a magnetic digital stirrer set at 400 rpm for 2 hours at room temperature to evaporate the organic solvent. Particles were collected by centrifugation at 1000 xg for 10 minuntes (Eppendorf centrifuge 5864 R, Eppendorf North America, Hauppauge, NY) and washed three times by discarding the supernatant, and resuspending in 30 mL of Ultrapure distilled water (Invitrogen, Waltham, MA). After the last washing step, the particle pellet was frozen in at -80°C and then dried under vacuum using a lyophilizer (Labconco Free zone 4.5L–105 °C, Labconco, Kansas City, MO). Supplemental figure 3 has a graphical depiction of protocol for preparing SARM-loaded PLGA microparticles.

### Characterization of SARM-loaded Microparticles: Particle Size and Surface Morphology

The particles size and surface morphology for each formulation were characterized using a Hitachi S-4800 scanning electron microscope (SEM, Hitachi High Technologies, Ontario, Canada) as described previously (*28, 75*). Briefly, a small amount of the particles was gently spread onto carbon double-adhesive tape mounted on an aluminum stub. To make the particles electrically conductive, an argon beam K550 sputter coater (Emitech Ltd., Kent, U.K) was used to coat the particles with gold (Au) and palladium (Pd), and SEM photomicrographs were taken at 5.0 kV accelerating voltage. The diameter of microparticles (n=100 per formulation) were measured using ImageJ software (NIH, Bethesda, MA). To further obtain a metric for the uniformity of particle sizes within each formulation, a span analysis was performed where the diameter below which 90% (D_90_) of the particles fell was subtracted from the diameter below which 10% of the particles fell (D_10_) and the result was divided by the diameter below which 50% of the particles fell (D_50_). A lower span value denotes a more homogenous particle size distribution.

### Differential Scanning Calorimetry (DSC)

DSC thermograms were obtained using a TA DSC instrument (TA Instruments model Q20, New Castle, DE) coupled with a refrigerated cooling system (RCS90). Each sample (3-5 mg) was weighed and transferred to a Tzero aluminum pan, then covered and compressed with a Tzero lid. Pure dry nitrogen purge gas was set at 20 psi pressure and 40 mL/min flow rate. The scanning temperature for all samples was set in the range of 0 – 175 ^°^C at a 10 ^°^C/min heating rate.

### Powder X-ray Diffraction (PXRD)

To obtain powder X-ray diffraction patterns a Siemens D5000 diffractometer was used. The X-ray source was composed of Cu K_α_ X-rays with λ =1.51418Å. The diffractograms of the samples were recorded in the range 5° to 50° at 2Θ values using a step size of 0.02° and a dwell time of 0.5 sseconds.

### Drug Loading, Encapsulation Efficiency, and Yield Percentage

To measure the drug loading (DL) and encapsulation efficiency (EE), SARM-loaded PLGA microparticles (2-5 mg) were dissolved in acetonitrile and 10- and 100-fold dilutions of that solution were made using 50:50 acetonitrile:water. SARM concentration was determined using HPLC-UV. Drug loading, encapsulation efficiency and yield percentages of microparticles were determined using equations 1, 2, and 3, respectively.

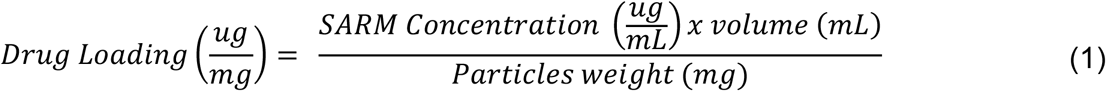

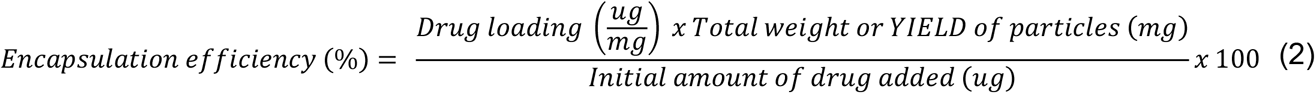

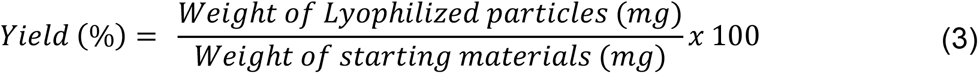

### In vitro Release Kinetics

To study in vitro release kinetics of SARM from the 2 microparticles formulations (F1 and F2), 5 mg of the SARM-loaded microparticles were suspended in 5 mL of release medium (0.4% v/v polysorbate 80 in 1 M PBS, pH = 7.4 release medium) and mixed at 300 rpm at 37^°^C. Tubes were centrifuged at 4500xg for 5 minuntes, 1 mL sample was withdrawn to be analyzed by HPLC-UV. Microparticles were resuspended in 5 mL of release medium after each withdrawal.

### In vivo Release Kinetics

For in vivo release kinetics, blood was collected via sub-mandibular blood draws and transferred to pre-heparinized tubes while the animal was anesthetized with 100-150 µL ketamine/xylazine (87.5/12.5 mg/kg, i.p.). Blood samples were centrifuged at 5000 xg and the supernatant (plasma) was separated and stored in -80°C freezer until ready to be processed and analyzed by HPLC-UV.

### Toxicity

To ensure the overall health of the mice following SARM-loaded microparticle administration, body weights were assessed throughout the 4 weeks of behavior testing. Animals were weighed twice per week for the duration of the experiment. To test for toxicity, we analyzed cardiac and liver toxicity after treatment with SARM-loaded microparticles. After the completion of behavior testing (end of week 4), animals were euthanized, and blood was collected via a cardiac blood draw. Blood was placed in 3.0 mL serum blood collection tubes (BD vacutainer, Franklin Lakes, NJ, USA) and allowed to clot for 30 minutes. The tubes were centrifuged at 1000 xg for 10 minutes at room temperature, serum was collected, and frozen. Frozen serum samples were sent to IDEXX (Columbia, MO, USA) for analysis of cardiac and liver toxicity panels. These panels detected serum levels of creatine kinase, alkaline phosphatase, aspartate transaminase, alanine aminotransferase, total bilirubin, albumin, and total protein. Immediately after collection of blood, the heart was collected and fixed in 10% neutral buffered formalin and routinely dehydrated through a series of ethanol and xylene baths, paraffin-embedded and sectioned at ∼4 µm onto glass slides for H&E staining. Stained tissue sections were given to a boarded American Academy of Veterinary Pathologist (ACVP) for examination.

### Statistical Analysis

All data is presented as mean±SEM, except for microparticle characterization and MWT values in supplemental table 1. Statistical analyses were performed on SPSS Version 25.0 (SPPS Inc. Chicago, IL) and GraphPad Prism Version 7.00 (GraphPad Software, La Jolla, CA). For MWT and paw mechanical sensitivity data, a repeated measures ANOVA containing the data points between day 1 and day 28 with baseline as a covariate was used to determine differences between groups. When appropriate, a Tukey’s post hoc test was used to determine group differences. For MWT and paw mechanical sensitivity data sets, Mauchly’s test of sphericity was implemented. If sphericity of data was not assumed, a Huynh-Feldt adjustment was utilized. However, in all data sets where a Huynh-Feldt was utilized the significance of the results did not change. Therefore, for clarity, we report the uncorrected F and p values in the results section. Effect sizes for MWT and paw mechanical sensitivity data is reported as Cohen’s *f* where 0.1 indicates small effect, 0.25 indicates medium effect, and 0.4 indicates large effect (*76*). Area under the curve (AUC) was calculated for MWT data by summing the change scores from D1 for each subsequent behavioral testing time point (D3-D28); zero represents no recovery of hyperalgesia. A student’s unpaired t-test (experiments 1+5) or one way ANOVA with Tukey’s post hoc test (experiment 4) was utilized to compare AUC between groups. An unpaired, student’s t-test was used to compare the average particle diameter size between SARM-loaded microparticle formulations. For serum cardiac and liver toxicant levels, an unpaired student’s t-test, with a Bonferroni correction for multiple comparisons, was used to determine differences between groups. For CPP data, a paired student’s t-test was used to compare time spent in SARM paired chamber between baseline and post drug pairing. For all data sets, we did not analyze the effect of sex as we were not powered to detect sex differences. However, we do report MWT means and standard deviations disaggregated by sex in Supplemental Table 1.

## List of Supplementary Materials

Supplemental Methods

Supplemental Figures 1-3

Supplemental Table 1

## Acknowledgements

The authors would like to thank Sudartip Areecheewakul for assisting with SEM photomicrographs of the microparticle formulations.

## Authors Contributions

Conceptualization: JBL, KAS, DSN, AKS

Methodology: JBL, KAS, DSN, AKS, ANP, SS

Data Collection: JBL, DSN, DKM, ANP, LR, AM, NG, PP

Funding acquisition: JBL, KAS, AKS

Project administration: JBL, KAS, DSN, AKS

Supervision: KAS, AKS

Writing – original draft: JBL, DSN

Writing – review & editing: JBL, KAS, DSN, AKS, DKM, SS

## Data and materials availability

Available upon request

