## Supplemental Files for "Selective Androgen Receptor Modulator Microparticle Formulation Reverses Muscle Hyperalgesia in Mouse Model of Widespread Muscle Pain"

### Supplemental Materials

### Supplemental Methods

#### High-performance Liquid Chromatography (HPLC-UV)

SARM was quantified in both aqueous solution and after extraction from mouse plasma using HPLC-UV coupled with a diode array detector (Agilent Infinity 1100, Santa Clara, CA). A reverse phase C18 column (5 µm pore size, 4.6 mm i.d.×150 mm) (Waters, Milford, MA, USA) was used for analysis. The mobile phase was a mixture of acetonitrile (0.1% trifluoro acetic acid):water (0.1% trifluoro acetic acid)(50:50 v/v) in an isocratic elution mode. The flow rate was set at 1 mL/min and the injection volume was set at 50 µL at room temperature. SARM was detected at 250 nm wavelength. (Supplemental Fig. 3A). A representative chromatogram of SARM in aqueous solution is depicted in (Supplemental Fig. 3B). A stock solution of SARM was prepared in pure ethanol at a concentration of 1 mg/mL and serial dilutions were prepared using acetonitrile:water (50:50 v/v) to construct the calibration curve in the range of 0.1–50 µg/mL (Supplemental Fig. 3C). A linear regression equation was fit to the calibration standards.

To quantify SARM in mice plasma samples, we constructed a calibration curve in plasma by spiking plasma (90 µL; Na heparin, mouse BALB/C plasma, Innovative research, Novi, Michigan) with 10 µL of SARM stock solutions (dissolved in ethanol in the range of 1 – 500 µg/mL) to result in a SARM concentration calibration range of 0.1 – 50 µg/mL (Supplemental Fig. 3D). Ivacaftor was selected as internal standard and 10 µL of a 10 µg/mL solution was spiked to the blank plasma samples for a (1 µg/mL) final concentration.

The SARM plasma calibration standards and plasma samples collected from mice from the pharmacokinetics experiment were extracted using an acetonitrile protein precipitation technique. Briefly, 1 mL acetonitrile was added to 100 µL of either the plasma calibration standards or the collected mice plasma samples (spiked with ivacaftor internal standard - 10 µL of 10 µg/mL solution), the mixture was vortexed for 5 min then incubated on ice for 10 min to allow time for plasma proteins to precipitate. Samples were centrifuged at high speed 10,000 xg for 5 min at 4°C and supernatant was then transferred to glass tubes and evaporated under a light stream of nitrogen. The residue was reconstituted in 100 μL 50:50 (v/v) acetonitrile:water, vortexed and centrifuged at 14,000 xg then 50 μL of the supernatant was analyzed by HPLC-UV.

### Supplemental Figures

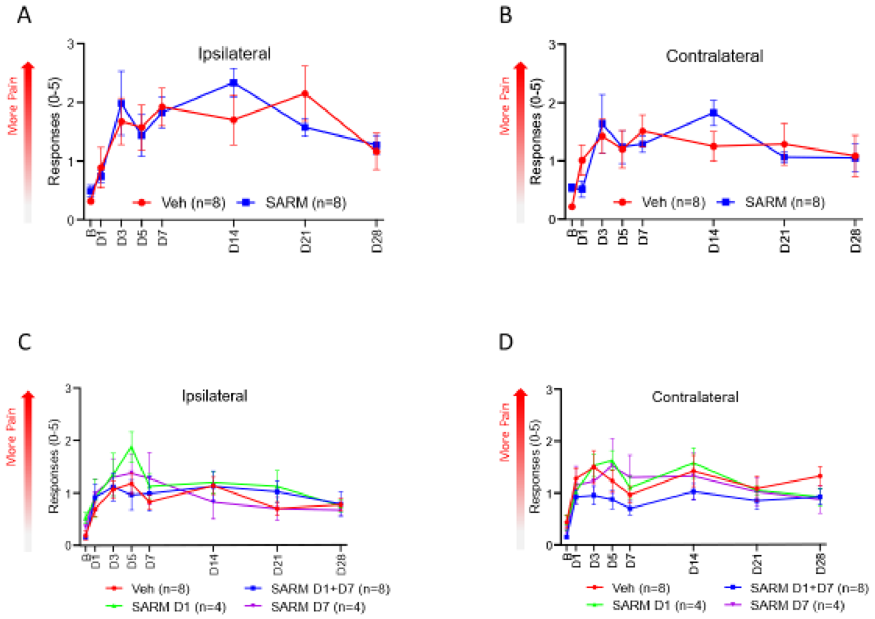


Supplemental Figure 1

**Daily SARM admnistration or SARM-loaded PLGA microparticles do not reverse mechanical hypersensitivity**. (A+B) Daily SARM administration was unable to alleviate paw hypersensitivity measured at the ipsilateral and contralateral hind paw when compared with vehicle treated animals. (C+D) SARM-loaded PLGA microparticles were unable to alleviate paw hypersensitivity on both the ipsilateral and contralateral hind paw when compared with vehicle treated animals. D=day, B=baseline, Veh=vehicle; Data are mean±SEM

- SARM-microparticles preparation: Single (O/W) emulsion solvent evaporation technique

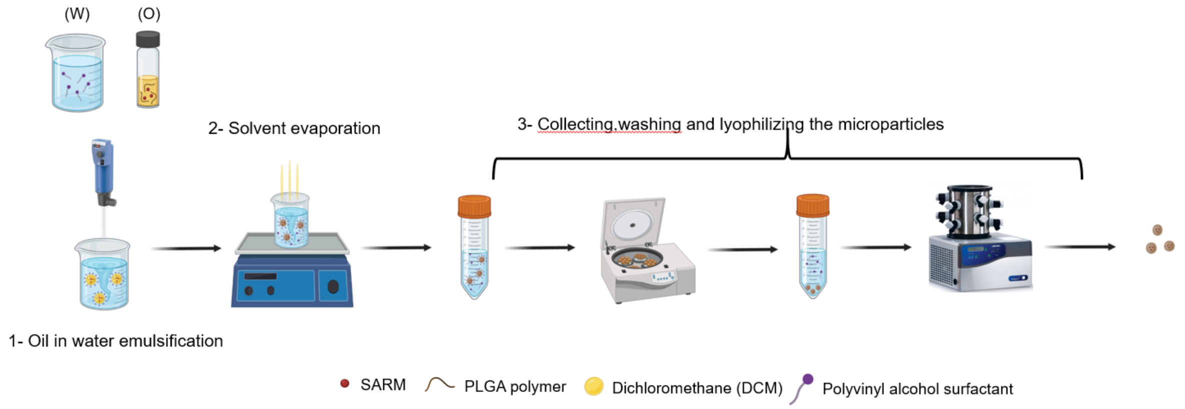


Supplemental Figure 2

Graphical Depiction of generation of SARM-loaded PLGA microparticles. Made in Bioredner. W=water, O=oil, PLGA=Poly (lactic-co-glycolic acid)

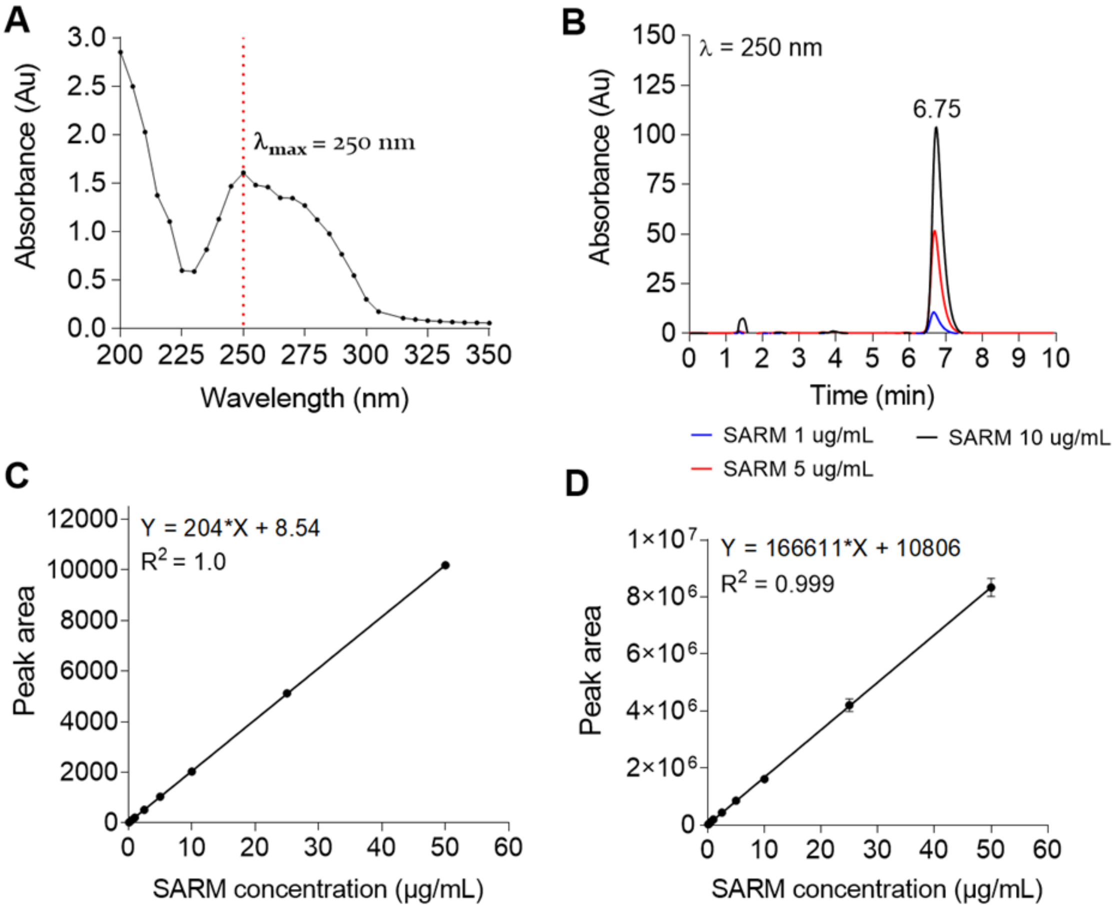


Supplemental Figure 3

(A) Ultra-violet (UV) absorbance spectrum of SARM showing maximum absorbance at 250 nm ($\lambda_{max}$ = 250 nm). (B) Representative HPLC-UV chromatogram of SARM in aqueous samples (50:50 acetonitrile:water, v/v) at concentrations of 1, 5, and 10 µg/mL. (C) HPLC-UV calibration curve of SARM in aqueous samples (acetonitrile:water, 50:50, v/v), showing good linearity in the range of 0.1 – 50 µg/mL ($R^2$ = 1.0). (D) HPLC-UV calibration curve of SARM extracted from plasma samples showing good linearity in the range of 0.1 – 50 µg/mL ($R^2$ = 0.999).

### Supplemental Table

| | | | | Ipsilateral | | | | |
|---|---|---|---|---|---|---|---|---|
| Testing Day | B | D1 | D3 | D5 | D7 | D14 | D21 | D28 |
| **Experiment 1 (Daily SARM)** | | | | | | | | |
| Male Veh | 1623.1±115.8 | 1083.9±75.7 | 1018.7±70.6 | 1120.9±159.1 | 1161.5±65.6 | 1097.2±77.9 | 1055.3±22.5 | 1435.1±133.1 |
| Female Veh | 1609.25±115.0 | 1045±164.4 | 1078.75±136.5 | 1125.5±227.3 | 1424.5±299.8 | 1334.3±289.5 | 1209.7±230.8 | 1532.6±107.1 |
| Male SARM | 1613.1±142.6 | 1122.7±28.8 | 1185.6±25.0 | 1278.1±176.4 | 1644.5±99.6 | 1572.1±141.9 | 1537.6±40.3 | 1575.2±136.0 |
| Female SARM | 1626.1±88.5 | 1079.6±62.2 | 1099.1±25.7 | 1237.3±105.6 | 1278.0±148.2 | 1339.5±189.2 | 1539.5±73.3 | 1584.2±60.3 |
| **Experiment 4 (SARM Microparticles)** | | | | | | | | |
| Male Veh | 1584.1±82.3 | 932±56.6 | 1023.5±122.6 | 1002.7±52.5 | 1112.2±206.5 | 1051.7±53.2 | 1062.2±50.5 | 1062.4±48.4 |
| Female Veh | 1569.5±43 | 1006±166.4 | 985.1±89.6 | 1025±54.4 | 1120.5±201.4 | 1139.1±209.4 | 1111.5±200.8 | 1125.5±82.3 |
| Male SARM D1+D7 | 1595.3±39.8 | 967±54.4 | 1036.3±114.8 | 1104.5±100.3 | 1299.5±280.2 | 1359.5±225.2 | 1402.1±76.8 | 1595.2±44.9 |
| Female SARM D1+D7 | 1588.4±27.4 | 1003.1±84.9 | 977.2±36.4 | 966.5±25.5 | 1162.4±278.8 | 1322.5±245.6 | 1399.1±223.8 | 1479.5±42.5 |
| Male SARM D1 | 1553.3±0.9 | 994.1±7.7 | 1063.6±44.7 | 1183.8±202.4 | 1213±120.2 | 1213.6±82.9 | 1126.1±69.1 | 1300.8±86.5 |
| Female SARM D1 | 1586±42.4 | 959.1±176.1 | 1012.8±55.3 | 1055.5±8.7 | 1111.5±169.9 | 1135.8±10.1 | 1192.6±11.7 | 1297.3±20.7 |
| Male SARM D7 | 1691.3±1.8 | 1133.3±23.5 | 1062.3±62.6 | 1154.1±154.3 | 1174±59.8 | 1319.1±153.9 | 1381.1±75.6 | 1317.5±53 |
| Female SARM D7 | 1589.5±34.1 | 1000.6±106.1 | 1058.3±17.9 | 1080.3±158.8 | 1108.3±1.8 | 1175.3±101.3 | 1164.5±22.3 | 1357.5±112.4 |
| **Experiment 5 (Flutamide Pellets)** | | | | | | | | |
| Male Veh | 1652±11.7 | 1023.1±77.1 | 1083.2±60.5 | 1215.5±83.1 | 1330.5±57.2 | 1522.1±91.8 | 1556.1±79.5 | 1583.4±48.9 |
| Female Veh | 1625.5±58.1 | 1022.7±145.2 | 1026.5±36.2 | 1103±49.8 | 1234.7±87.4 | 1389.1±46.8 | 1440.3±29.3 | 1490.5±106.6 |
| Male Flutamide | 1666.4±55.9 | 978.5±97.8 | 1034.6±47.9 | 1028±86.2 | 1046±14.1 | 1044.1±141.6 | 1183.9±103.5 | 1174.9±147.2 |
| Female Flutamide | 1576.1±57.6 | 990.6±43.2 | 994±120.7 | 1020.6±66.5 | 1054.7±45.1 | 1064.7±59.7 | 1110.9±49.6 | 1105.5±72.5 |
| | | | | Contralateral | | | | |
| Testing Day | B | D1 | D3 | D5 | D7 | D14 | D21 | D28 |
| **Experiment 1 (Daily SARM)** | | | | | | | | |
| Male Veh | 1666.1±177.7 | 1079.1±78.3 | 1057.0±49.0 | 1122.0±144.0 | 1125.0±107.4 | 1177.5±214.7 | 1098.5±136.8 | 1366.4±110.5 |
| Female Veh | 1590.1±73.9 | 1083.5±111.5 | 1120.8±127.5 | 1091.9±159.8 | 1360.8±287.2 | 1297.1±262.2 | 1237.7±193.3 | 1446.3±135.9 |
| Male SARM | 1622.2±142.9 | 1107.6±89.8 | 1150.8±46.2 | 1138.1±110.1 | 1530.7±151.7 | 1485.8±136.8 | 1494.9±72.7 | 1512.7±2.6 |
| Female SARM | 1615.6±152.9 | 1145.4±69.2 | 1114.1±73.4 | 1199.8±66.8 | 1232.2±180.5 | 1326.5±212.2 | 1575.1±64.5 | 1507.9±75.8 |
| **Experiment 4 (SARM Microparticles)** | | | | | | | | |
| Male Veh | 1677.5±52.5 | 1013.6±30.7 | 1051.8±41.1 | 1079.7±60.6 | 1089.4±89.2 | 1078.4±52.5 | 1120±117.4 | 1113.1±113.2 |
| Female Veh | 1656.5±60.7 | 1016±105.2 | 1023.1±38.8 | 1028±69.6 | 1226±183.6 | 1154.4±285.5 | 1141.1±248.2 | 1127.3±52.8 |
| Male SARM D1+D7 | 1680.5±50.3 | 1053.7±89.3 | 1106.8±71.8 | 1142.5±131.6 | 1198.4±201.8 | 1357.7±198.5 | 1537.1±67.5 | 1609±32.9 |
| Female SARM D1+D7 | 1637.5±49.7 | 991.9±43.5 | 948.1±54.1 | 1049.5±83.2 | 1127.2±92.2 | 1372.5±294.9 | 1413.5±176.9 | 1504.1±92.3 |
| Male SARM D1 | 1637.3±24.5 | 1127.3±103.7 | 1024.3±3.7 | 1119.3±230 | 1145±57.5 | 1234.6±54.2 | 1100±71.1 | 1358.6±185.2 |
| Female SARM D1 | 1588.5±51.1 | 1035.3±1.4 | 945.8±24.7 | 993±70.7 | 1104.8±137.4 | 1199±32.1 | 1190±57.9 | 1262.8±130.8 |
| Male SARM D7 | 1702±44.3 | 1052.5±77.1 | 1042.3±51.3 | 1121.6±90 | 1223.5±139.3 | 1265.3±16 | 1357±7.5 | 1434.3±123.9 |
| Female SARM D7 | 1589.5±34.1 | 1000.6±106.1 | 1058.3±17.9 | 1080.3±158.8 | 1108.3±1.8 | 1175.3±101.3 | 1164.5±22.3 | 1357.5±112.4 |
| **Experiment 5 (Flutamide Pellets)** | | | | | | | | |
| Male Veh | 1719±62.6 | 1039.4±36.3 | 1057.8±55 | 1318.1±114.6 | 1316.2±86.3 | 1499±88.5 | 1615.9±29.3 | 1564.4±81.4 |
| Female Veh | 1671.4±47.2 | 1075.4±85.4 | 1081.8±85.7 | 1185.5±140.9 | 1296.3±169.8 | 1339.4±107.3 | 1452.3±70.9 | 1496.6±64 |
| Male Flutamide | 1722.5±42.7 | 990.5±50.6 | 1025.9±65.3 | 1062.7±19.8 | 1061.1±16.8 | 1058.5±57.3 | 1136.5±94.6 | 1134.7±176.8 |
| Female Flutamide | 1604.5±103.3 | 985.1±31.9 | 1014.1±27.3 | 1028.1±50.9 | 1058.8±51.7 | 974±100.9 | 1068±113.2 | 1075.9±50.9 |

Supplemental Table 1

Muscle withdrawal threshold data for each experiment disaggregated by sex. B=Baseline, D=Day, Veh=Vehicle; Data is mean±SD.
